# Grey matter biomarker identification in Schizophrenia: detecting regional alterations and their underlying substrates

**DOI:** 10.1101/357954

**Authors:** V. Chatzi, R.P. Teixeira, J. Shawe-Taylor, A. Altmann, O. O’Daly, D. Christiaens, J. Schrouff, J.-D. Tournier

## Abstract

State-of-the-art approaches in Schizophrenia research investigate neuroanatomical biomarkers using structural Magnetic Resonance Imaging. However, current models are 1) voxel-wise, 2) difficult to interpret in biologically meaningful ways, and 3) difficult to replicate across studies. Here, we propose a machine learning framework that enables the identification of sparse, region-wise grey matter neuroanatomical biomarkers and their underlying biological substrates by integrating well-established statistical and machine learning approaches. We address the computational issues associated with application of machine learning on structural MRI data in Schizophrenia, as discussed in recent reviews, while promoting transparent science using widely available data and software. In this work, a cohort of patients with Schizophrenia and healthy controls was used. It was found that the cortical thickness in left pars orbitalis seems to be the most reliable measure for distinguishing patients with Schizophrenia from healthy controls.

**Highlights:**

- We present a sparse machine learning framework to identify biologically meaningful neuroanatomical biomarkers for Schizophrenia
- Our framework addresses methodological pitfalls associated with application of machine learning on structural MRI data in Schizophrenia raised by several recent reviews
- Our pipeline is easy to replicate using widely available software packages
- The presented framework is geared towards identification of specific changes in brain regions that relate directly to the pathology rather than classification per se

## 1 Introduction

Schizophrenia is a mental health disorder that can emerge at late adolescence/early adulthood and last for a lifetime. Its onset is often followed by deterioration of cognition, perception and social behaviour (Tamminga & Medoff, 2000). Depending on the magnitude of these deficits as well as the severity and frequency of the clinical symptoms, everyday-life coping is frequently impaired and followed by social withdrawal, unemployment, suicidal thoughts or behaviour (Hooley, 2010; Hor & Taylor, 2010). These complex personal and social issues are well known to the research community and have triggered an increased interest for developing methods that facilitate detection of early signs of the disorder, which in turn facilitates interventions, disease management and therapy provision before the expression of the full onset of the symptoms (Modinos & McGuire, 2015).

Structural Magnetic Resonance Imaging (MRI) has been widely used to study these early signs *in vivo*. Using this non-invasive technique, a growing literature in anatomical brain abnormalities associated with Schizophrenia has emerged (Fusar-Poli et al., 2012). Of particular interest is the potential utility of structural MRI data for identifying neuroanatomical markers that are reproducible across studies and translatable to the clinical setting (Woo et al., 2017).

Previous studies have shown that it is possible to classify patients with Schizophrenia from healthy controls using structural MRI data (Orru et al 2012). However, 1) these frameworks often use voxel-wise, whole brain input features, leading to high-dimensional input feature spaces which is problematic given their small sample size (Varoquaux, in press). Such problems are ill-posed and without the use of regularisation approaches to ensure a unique solution of the model, these models have increased computational complexity and a tendency to overfit. 2) In addition, they use voxel-wise input features known as grey matter density maps to quantify morphological alterations. Morphological alterations can occur due to several factors including changes in volume, cortical thickness or gyrification (Mechelli et al., 2005). As a result, grey matter density maps, as well as the resulting output weight maps are able to localise grey matter alterations but not their underlying substrates. They are therefore poorly interpretable in terms of the biological nature of the grey matter changes. 3) Lastly, previous frameworks’ pipelines as well as their findings are poorly generalisable. As a result, reported accuracy values for predicting patients with Schizophrenia from healthy controls range between 60-95% (Arbabshirani et al., 2017).

To address these limitations, in the present work 1) we extract low dimensional, region-wise brain measures, and introduce stability selection to identify the most stable features for the classification; 2) we generate region-wise output maps that reflect the substrates of the region-wise grey matter alterations to boost interpretability and facilitate understanding of the pathophysiology of the disorder; 3) we use well-established, widely available statistical and machine learning packages to build a robust and generalisable framework, and a publicly available dataset to enable reproducibility of the presented pipeline and research findings.

Our framework focuses on identifying grey matter regions which are meaningful for the understanding of the psychopathology of Schizophrenia by addressing previously reported methodological limitations of statistical and machine learning frameworks. To the authors’ knowledge, this is the first study in Schizophrenia research to address these limitations within a unified machine learning framework, and to identify biologically meaningful anatomical alterations in critical regions for Schizophrenia. This pipeline has the potential to impact on future studies that investigate neuroanatomical biomarkers within niche groups of the disorder such as First Episode Psychosis, At-Risk-Mental-State, treatment response or treatment resistance, and for drug development to target the identified regions.

## 2 Methods

### 2.1 Materials

#### 2.1.1 Participants

In this study, 146 participants were used from the publicly available dataset released through the Centre of Biomedical Research Excellence (COBRE) consortium (Çetin et al., 2014; Aine et al., 2017). Participants were 18-65 years old (mean age=37.42±12.75), 112 were male and 34 were female. Sixty-three were patients with Schizophrenia (male=48, female=15, age=37.19±13.80), diagnosed with the structured clinical interview for DSM-IV axis I disorders (SCID) and were medicated and 83 were healthy controls (male=64, female=19, age= 37.59±11.98).

#### 2.1.2 Image acquisition

Images were acquired on a Siemens 3T Trio TIM scanner using a 12-channel head coil. Structural MRI data consist of high-resolution T1-weighted MPRAGE images (TR: 2530 ms, TI: 900ms, flip angle: 7°, FOV: 256×256 mm, slab thickness: 176mm, 1mm isotropic resolution).

### 2.2 Image pre-processing

Pre-processing of the structural images was performed using Freesurfer version 5.3.0 (Fischl, 2012). The full Freesurfer pre-processing pipeline was performed, including full surface reconstruction and volumetric segmentation (Dale et al., 1999; Dale and Sereno, 1993; Fischl and Dale, 2000; Fischl et al., 2001; Fischl et al., 2002; Fischl et al., 2004a; Fischl et al., 1999a; Fischl et al., 1999b; Fischl et al., 2004b; Han et al., 2006; Jovicich et al., 2006; Segonne et al., 2004, Yeo et al., 2010a; Yeo et al., 2010b). The used steps included motion correction, skull-stripping (Segonne et al. 2004), Talairach transformation (Collins et al.,1994), subcortical segmentation (Fischl et al., 2002; Fischl et al., 2004a, Segonne et al., 2003), intensity normalization (Sled et al., 1998), grey matter / white matter boundary tessellation, topology correction (Fischl et al., 2001; Segonne et al., 2007; Segonne et al., 2005), surface deformation (Dale et al., 1999; Dale and Sereno, 1993; Fischl and Dale, 2000), surface inflation (Fischl et al., 1999a), registration to a spherical atlas (Fischl et al., 1999b), parcellation (Desikan et al., 2006; Fischl et al., 2004b), and generation of surface and subcortical statistics were performed.

### 2.3 Quality control

All extracted masks, including the extracted brain, the white matter boundaries, the pial surface boundaries, along with the subcortical segmentation were overlaid onto the T1-weighted images and their quality was visually checked across all slices and planes using freeview (https://surfer.nmr.mgh.harvard.edu/fswiki/FreeviewGuide/FreeviewGeneralUsage/FreeviewQuickStart). All surfaces were visually checked using tksurfer (https://surfer.nmr.mgh.harvard.edu/fswiki/TkSurfer).

### 2.4 Feature extraction

After pre-processing and quality control, the resulting subcortical segmentation files and parcellation files that were created using the Desikan-Killiany atlas (Desikan et al., 2006) from each scan were used to form the initial feature matrix. In total, 68 cortical regions using the Desikan-Killiany atlas were identified, and 17 subcortical regions using the segmentation files that resulted from the Freesurfer pre-processing, namely: Cerebellum, Thalamus, Caudate, Putamen, Pallidum, Hippocampus, Amygdala, and Accumbens-area (Long et al., 2017) and Brain Stem. Following definition of the cortical and subcortical regions, the grey matter measures were extracted region-wise. Six cortical features namely cortical thickness, surface area, mean curvature, folding index, curving index, and cortical volume were extracted for each of the 68 cortical regions and 1 subcortical feature namely subcortical volume was extracted for each of the 17 regions (Figure 1). Therefore, the total feature set comprised of 425 features.

**Figure 1.**
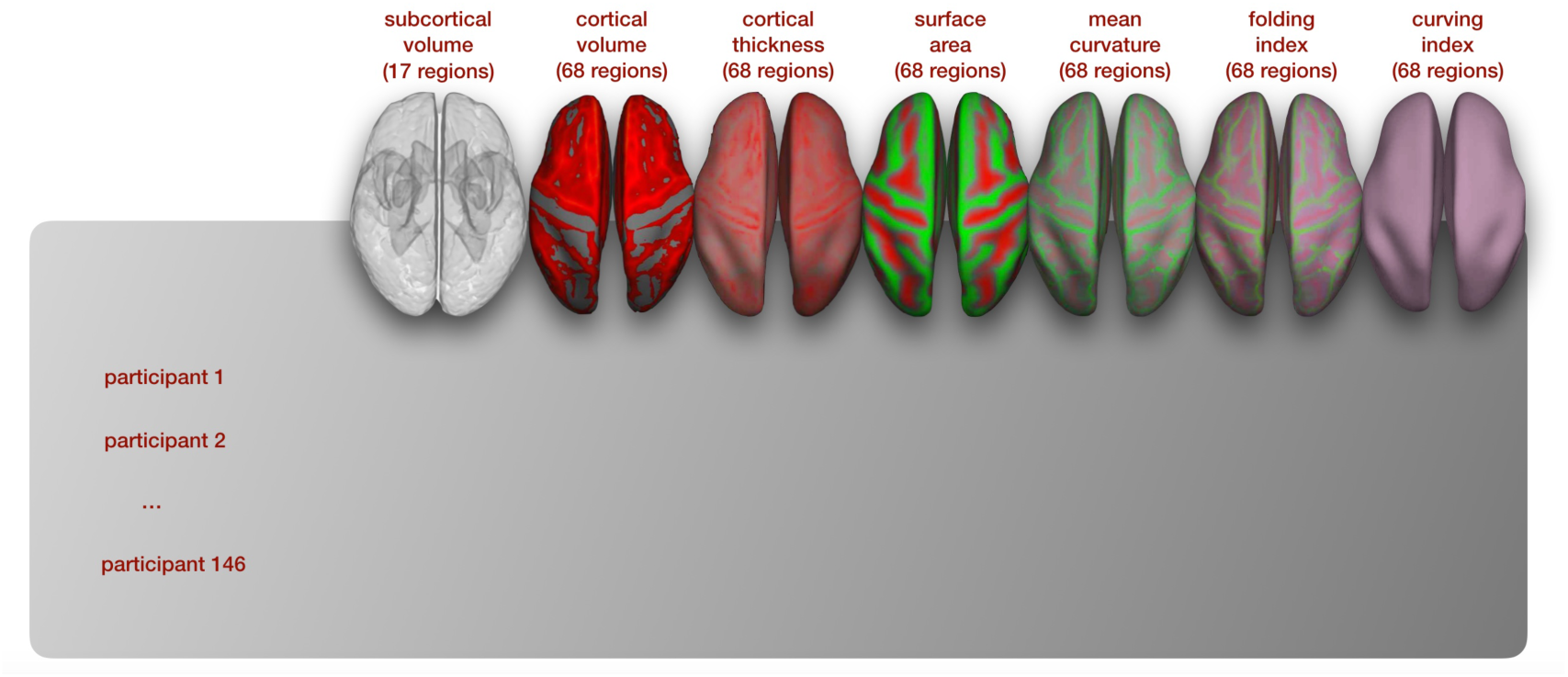
Extracted features: cortical thickness, surface area, mean curvature, folding index, curving index, and volume for 68 cortical regions, and volume for the 17 subcortical regions

**Figure 2.**
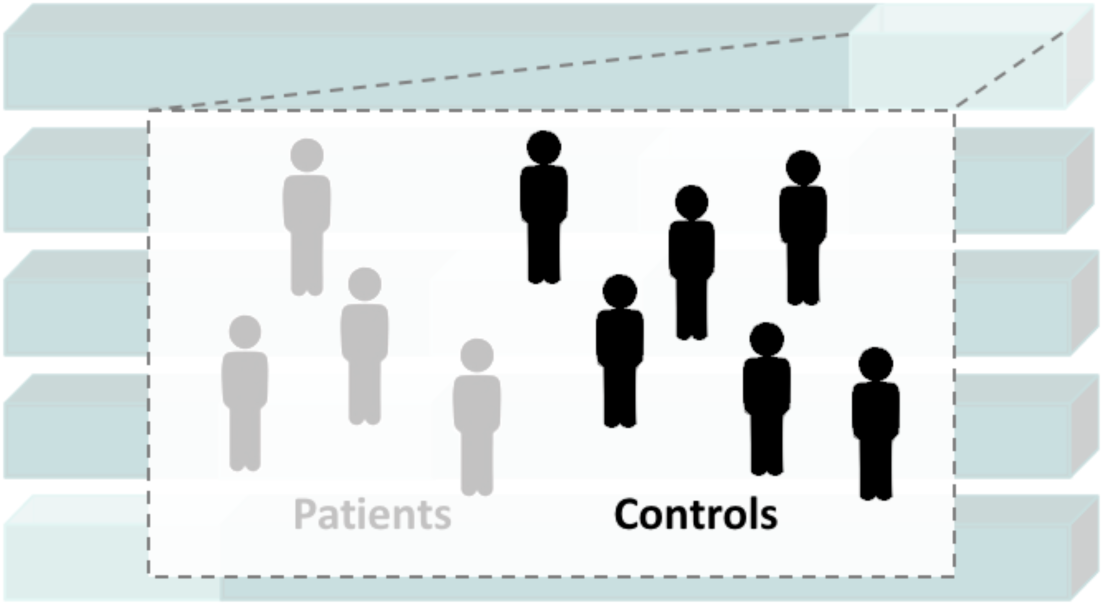
Each training and tests were split using stratification, maintaining at each split 40% patients and 60% controls.

### 2.5 Machine learning modelling

The main framework for the machine learning analysis consisted of two main steps, namely stability selection and SVM classification. The framework integrated holdout validation and stratification of the data and is illustrated in figure 3.

**Figure 3.**
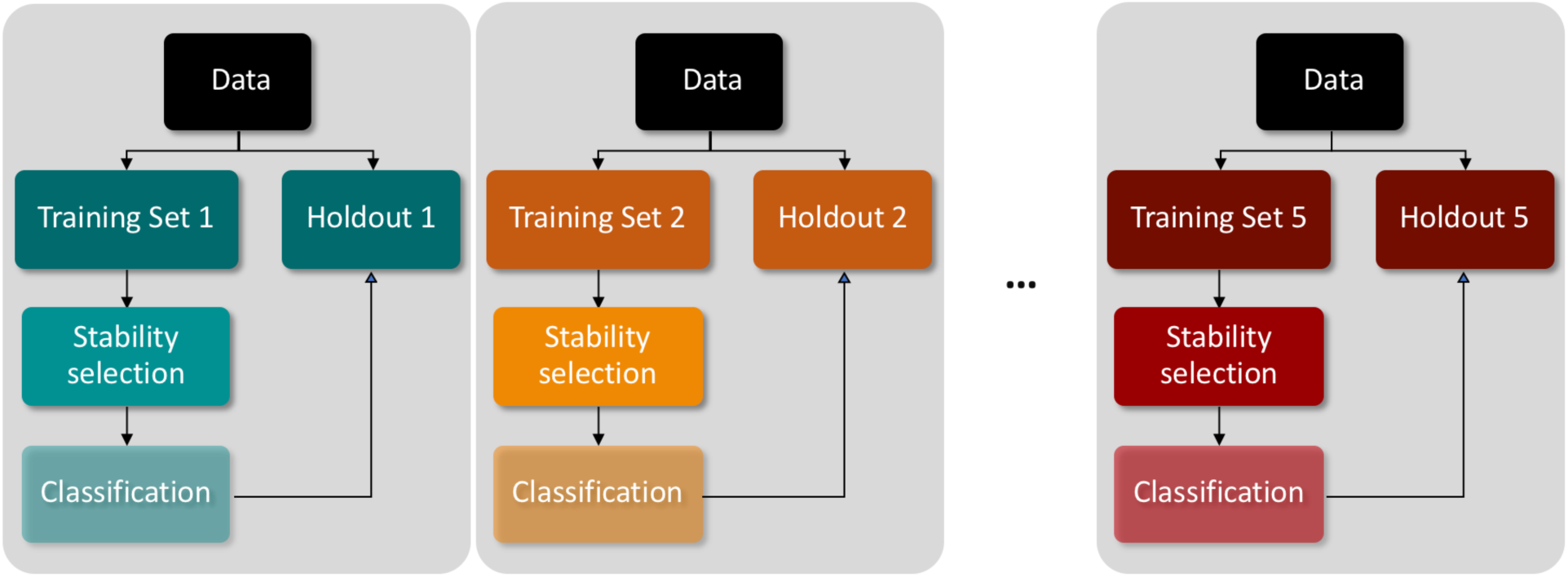
Data was split in training set and holdout (or test set) in five different ways. The training set was used for stability selection and classification and the test set for assessing generalisability of the trained classifier to unseen data.

#### 2.5.1 Stratification

The dataset was slightly imbalanced, with 83 healthy controls and 63 patients with Schizophrenia. Stratification enabled the proportional inclusion of participants from the patient group and the control group in each split of the data (Nichols, 2016). Specifically, each split of the data comprised of 40% patients with Schizophrenia and 60% healthy controls (Figure 2).

#### 2.5.2 Holdout validation

Five holdouts were defined (Figure 3). Each holdout set was generated with stratification and included 20% of the total dataset. Each subject was included only within one holdout. Therefore, each holdout was completely different from all other holdouts. Each holdout fold enabled the definition of 5 different training datasets and 5 different test sets. Each holdout served as a test set, and the remaining 80% of the data was used as training set for stability selection and training of the model. Importantly, holdout sets were only used to assess the generalizability of the estimated model after stability selection and classification training was performed on the training set.

#### 2.5.3 Stability selection

Stability selection was performed in R using the package stabs (Hofner & Hothorn, 2017; Hofner et al., 2015; Meinshausen & Buehlmann, 2010; Shah & Samworth, 2013). The stable features were selected using The Least Absolute Shrinkage and Selection Operator (LASSO, Tibshirani, 1996) glmnet. Features were assessed as ‘stable’ in the training set if they were selected in at least 60% of the times (Tibshirani, 1996).

#### 2.5.4 Support Vector Machines for binary classification

Following Stability selection, classification training was performed using Support Vector Machines (SVMs) (Cortes & Vapnik,1995; Cristianini & Shawe-Taylor, 2000) using the *fitcsvm* function from MatLab 2017b classification toolbox (The MathWorks, Natick, MA, USA). Support Vector Machines for classification, search for a linear separating hyperplane between the two classes in a feature space that is defined by the kernel. The optimal hyperplane is defined as the one with the largest distance from the nearest observations (the support vectors) from the two classes, known as the maximum-margin hyperplane and the width of the hyperplane characterizes the ability to separate confidently between the two classes. Two different kernel SVMs were explored for binary classification available in matlab: Linear and Quadratic (Table 1).

**Table 1.** kernel formulas used

| **Kernel function** | **formula** |
| --- | --- |
| *Linear* | $G(x_j, x_k) = x_j'x_k$ |
| *Quadratic* | $G(x_j, x_k) = (1 + x_j'x_k)^2$ |

#### 2.5.5 Hyperparameter optimisation

Hyperparameter optimisation for the linear and Quadratic SVMs were optimised using the function bayesopt for Bayesian optimisation (Martinez-Cantin, 2014) with a 5-fold cross validation in MatLab 2017b classification toolbox (The MathWorks, Natick, MA, USA).

#### 2.5.6 Model evaluation

Following training and optimization of the classifiers’ hyperparameters, each of the classifiers was applied to the holdout, to test generalizability of the fitted model to the unseen data. Subsequently the testing accuracy for each outer loop was calculated, followed by the class accuracies, f score and geometric mean.

## 3 Results

### 3.1 Stability selection

During stability selection, for training set 1 and 5 the cortical thickness of the left pars orbitalis (red) was selected (figure 4) while for folds 2, 3 and 4, no features were selected.

**Figure 4.**
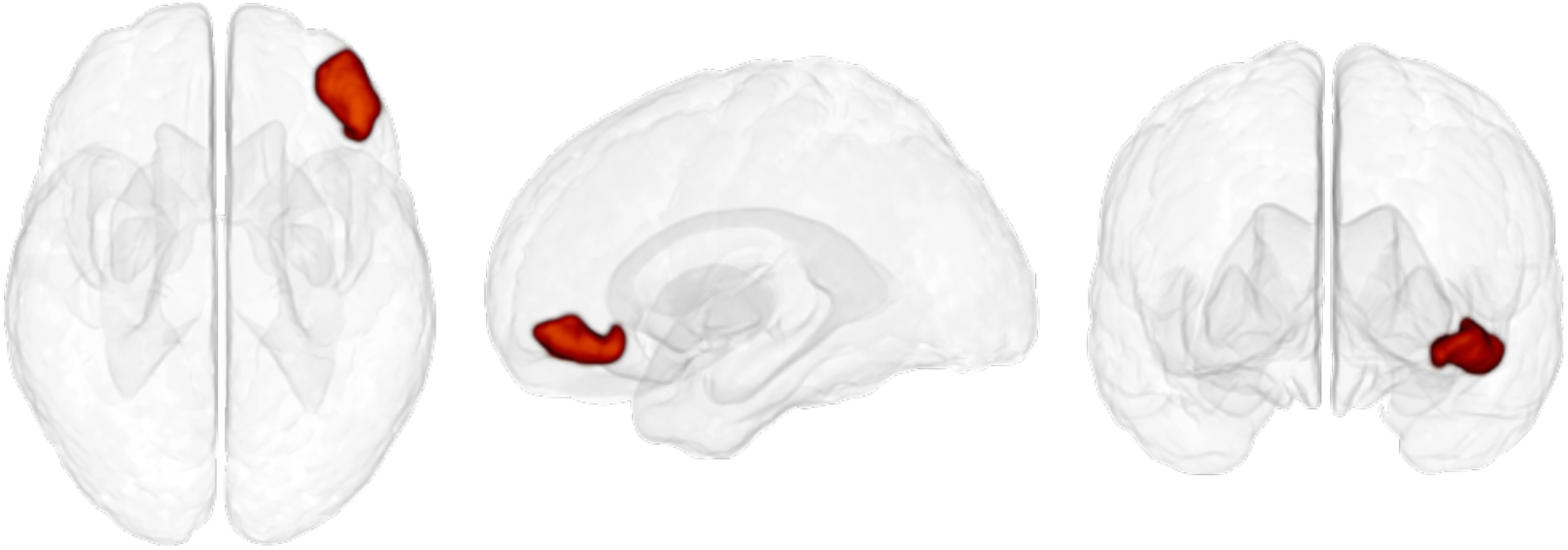
Stable feature.: cortical thickness of left pars orbitalis also known as Broca’s area.

### 3.2 Training, Model evaluation and generalisability assessment

Following stability selection, the training sets 1 and 5 were trained using the cortical thickness in the left pars orbitalis as input feature. Then the trained models were applied in the equivalent holdout set and the performance metrics for linear and quadratic SVMs on the test set were calculated. The averaged testing accuracy using the Linear model was 64.29% while for the Quadratic model was 67.86% (Table 2). Controls were predicted a bit better than patient group and overall the quadratic kernel SVM performed better compared to linear kernel SVM. Performance metrics were calculated accordingly (Table 2).

**Table 2.**
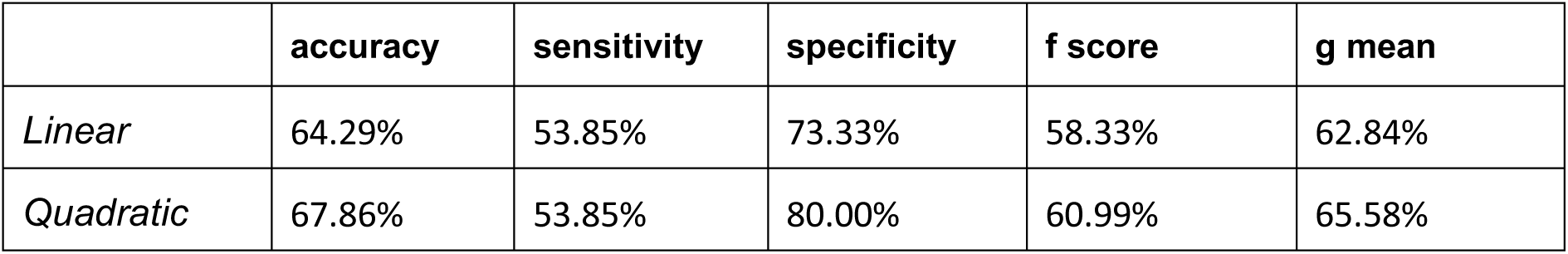
model comparison in test set, assessing generalisability

|  | accuracy | sensitivity | specificity | f score | g mean |
| --- | --- | --- | --- | --- | --- |
| *Linear* | 64.29% | 53.85% | 73.33% | 58.33% | 62.84% |
| *Quadratic* | 67.86% | 53.85% | 80.00% | 60.99% | 65.58% |

## 4 Discussion

In this work, we developed a data-driven pipeline to identify features implicated in the pathology, rather than for classification as such. Our conservative method for feature selection, enabled the identification of changes that are robust and generalisable, and the output of interest are the features identified. We achieved this by addressing computational issues and common pitfalls associated with application of machine learning on structural MRI data in Schizophrenia as discussed in recent reviews (Arbabshirani et al., 2017; Varoquaux, in press; Woo et al., 2017; Smith & Nichols, 2017) using well-established and widely available software.

First, we addressed the issue of high dimensionality of MR Images, which is problematic given the small sample size of neuroimaging studies. Previous frameworks have primarily relied on voxel-wise features, whereby each image voxel is considered a separate feature, resulting in enormously dimensional input for the classifier, on the order of several hundreds of thousands to millions of features (depending on image resolution, Monteiro et al., 2017). This input feature space is particularly problematic in Schizophrenia research where imaging datasets typically comprise a few dozen participants. Specifically, common datasets consist of a few, very high-dimensional feature vectors. The lack of dimensionality reduction methods to reduce this high-dimensional feature space results in a computational problem that is prone to overfitting thus leading to a model that generalizes poorly to unseen data. Even imposing different regularization can lead to the identification of different discriminative patterns for similar accuracies (Baldassarre et al., 2012; 2017). These problems with high dimensional features and few samples are also ill-posed with multiple solutions that all provide similar levels of accuracy. Therefore, reducing the high dimensionality of the imaging data is crucial to find the solution to the classification problem.

To address the high dimensionality of the data as opposed to the small sample size, we used feature extraction and stability selection methods. Specifically, we extracted from the structural MR Images, cortical grey matter measures using the Desikan-Killiany parcellation atlas and automatically segmented subcortical region-wise grey matter measures as provided by the FreeSurfer pre-processing pipeline. Although cortical volume, subcortical volume, and cortical thickness have been previously used for classification in Schizophrenia research, curvature, folding index, curving index and surface area have not previously been explored for this purpose. Although curvature, folding index, curving index and surface area did not seem to drive the classification in this cohort of chronic patients with Schizophrenia versus healthy controls, these measures have been shown to be significantly altered in niche cohorts within Schizophrenia research such as first episode psychosis and treatment response cohorts (Gur et al., 1999; Kuperberg et al., 2003; Palaniyappan et al, 2011; Palaniyappan et al, 2012; Palaniyappan et al., 2013; 2016; Wagstyl et al., 2016) and therefore we would expect that they could drive the classification in those cohorts.

Following extraction of low-dimensional region-wise grey matter measures, we identified the most informative for the classification features using well-established and freely available software for stability selection (Hofner & Hothorn, 2017; Hofner et al., 2015; Meinshausen & Buehlmann, 2010; Shah & Samworth, 2013). Stability selection enabled us to identify the most relevant and stable for the classification features. Out of the 425 features we were able to identify one stable feature. This feature was the cortical thickness of the left pars orbitalis. Alterations in left pars orbitalis or Broca’s area in Schizophrenia is a well-replicated finding including cerebral blood flow (McGuire et al., 1993) and cortical thickness (Viher et al 2018; Ehrlich et al.,2012; Selemon et al., 2003). Therefore, the incorporation of stability selection facilitated the interpretability of our findings.

The identification of the substrate of the alteration in left pars orbitalis, namely the cortical thickness as a single neuroimaging biomarker for Schizophrenia, is the most distinctive contribution of this work in comparison to previous classification frameworks. Although previous frameworks were capable of localizing patterns of morphological alterations, they were not able to identify the biological substrates of the resulting changes e.g., volume, cortical thickness or gyrification (Mechelli et al., 2005). The most commonly reported classification output amongst most previous machine learning frameworks is a voxel-wise map of inter-correlated voxel weights, showing the contribution of each voxel to the overall classification with respect to all other voxels. However, due to the inter-dependencies between voxels inherent in the model, no inference can be made either for an individual voxel independently of all others, or for the regions of interest (ROIs), or for the underlying substrate (e.g., volume, cortical thickness, gyrification) (Taylor et al. 2017). In this work we showed that it is possible to identify the most relevant feature that can serve as biomarkers to classify patients with Schizophrenia from healthy controls and draw inference on the contribution of these biomarkers to the overall classification.

Following stability selection, we introduced stratification during training of the three classifiers. To the authors knowledge this is the first study to introduce stratification within a machine learning framework in Schizophrenia. The use of stratification is crucial for Schizophrenia research where imaging datasets are often imbalanced and commonly one group of interest outnumbers the other. Most studies do not take into account this imbalance during training and testing of the classifier, leading to a number of issues including largely imbalanced proportions of the groups of interest within each split/fold or even absence of participants from both groups of interest within one or more splits/folds, which can then lead to biased findings. This issue was addressed with stratification, where the same proportion of the groups of interest was kept stable within and across all splits of the data. This way it was possible to keep representative proportions of each group of interest within each split of the data and measure the effect of interest between our groups.

We then performed model evaluation on the holdout set. The reason for reporting on the test set is because folds with very high accuracy at the training stage may not necessarily perform well on the unseen data. This process is essential to minimize the risk of reporting a model that is overfitting the data, and therefore maximize the possibility of achieving a good bias-variance trade-off and good generalizability while accounting for model complexity. This trade-off is often ignored, and the model with the highest training accuracy is typically reported. As a result, some studies report exceptionally high accuracies that cannot be replicated by others, as Arbabshirani et al. (2017) and Varoquaux (in press) discussed in their recent reviews. This is a matter of great concern, and it is important to integrate appropriate approaches to explore the generalizability of the developed models. To address this issue, we implemented two different models with different degrees of complexity, one with a linear kernel and one with a quadratic kernel, and subsequently compared them with respect to their testing accuracy. We focused on reporting the classification performance metrics in the testing set, in contrast to the majority of previous studies, to provide a realistic and conservative estimate of the potential of structural MRI data for biomarker identification in Schizophrenia.

Following addressing the computational limitations of previous frameworks, we were able to identify a biologically meaningful neuroimaging biomarker for Schizophrenia, with a testing accuracy that suggests that structural MRI may not be the most accurate modality to classify patients with Schizophrenia from healthy controls. In our future work we aim to investigate whether other MRI modalities may or may not be more powerful in their ability to classify this or other cohorts.

A limitation of the presented work is the lack of a large niche cohort such as individuals at risk for developing the disorder or first episode psychosis. We anticipate that this pipeline can be used for those cohorts to reveal biomarkers in these different stages of the disorder or its or trajectories over the life span. The pipeline can be also used for biomarker identification using other MRI modalities such as diffusion weighted MRI or functional MRI. We anticipate that this pipeline can be beneficial for multi-centre studies where data can be pre-processed separately in each neuroimaging site and merged together after pre-processing, as well as for multimodal frameworks where the extraction of low-dimensional, meaningful features from each modality will be essential for overcoming the demanding computational conundrums of such frameworks. We anticipate that this pipeline is a big step forward towards biomarker identification using widely available tools, while addressing the computational complexity of implementing machine learning on MRI data and that the additional interpretability attributed to this framework can facilitate our understanding of the dynamics of the physiopathology of the Schizophrenia as well as other psychiatric and neurodegenerative disorders.

## Acknowledgements

Vasiliki Chatzi is funded by the Department of Psychosis Studies at King’s College London. Data was downloaded from the COllaborative Informatics and Neuroimaging Suite Data Exchange tool (COINS; http://coins.mrn.org/dx) and data collection was performed at the Mind Research Network and funded by a Center of Biomedical Research Excellence (COBRE) grant 5P20RR021938/P20GM103472 from the NIH to Dr. Vince Calhoun. The main idea for this paper was developed from discussions with Peter Dayan, and the methods were optimised with help from Joao Monteiro, and Janaina Mourao-Miranda. Vasiliki Chatzi would like to thank Margaret King and Vince Calhoun for their continuous support throughout this project with respect to the dataset and the IoPPN mri support team for the technical support. Andre Altmann holds an MRC eMedLab Medical Bioinformatics Career Development Fellowship. This work was supported by the Medical Research Council [grant number MR/L016311/1].

## References

Aine, C.J., Bockholt, H.J., Bustillo, J.R. et al. Neuroinform (2017). Multimodal Neuroimaging in Schizophrenia: Description and Dissemination, 15: 343. https://doi.org/10.1007/s12021-017-9338-9

Arbabshirani, M.R., Plis, S., Sui, J., & Calhoun, V. D. (2017). Single subject prediction of brain disorders in neuroimaging: Promises and pitfalls. NeuroImage, 145, 137–165.

Baldassarre L, Pontil M, Mourão-Miranda J. (2017). Sparsity Is Better with Stability: Combining Accuracy and Stability for Model Selection in Brain Decoding. Frontiers in Neuroscience, 11:62. doi:10.3389/fnins.2017.00062

Baldassarre, L., Mourao-Miranda, J., Pontil, M. (2012). Structured sparsity models for brain decoding from fMRI data, Pattern Recognition in NeuroImaging (PRNI), International Workshop on, 5–8

Çetin, M., Christensen, F., Abbott, C., Stephen, J., Mayer, A., Cañive, J., Bustillo, J., Pearlson, G., and Calhoun V. D. (2014). “Thalamus and posterior temporal lobe show greater inter-network connectivity at rest and across sensory paradigms in Schizophrenia”, NeuroImage, vol. 97, pp. 117–126,

Collins, D.L., Neelin, P., Peters, T.M., and Evans, A.C. (1994). Automatic 3D Inter-Subject Registration of MR Volumetric Data in Standardized Talairach Space, Journal of Computer Assisted Tomography, 18(2) p192–205, 1994 PMID: 8126267; UI: 94172121

Dale, A.M., Fischl, B., Sereno, M.I., 1999. Cortical surface-based analysis. I. Segmentation and surface reconstruction. NeuroImage 9, 179–194.

Desikan, R.S., Segonne, F., Fischl, B., Quinn, B.T., Dickerson, B.C., Blacker, D., Buckner, R.L., Dale, A.M., Maguire, R.P., Hyman, B.T., Albert, M.S., Killiany, R.J., 2006. An automated labeling system for subdividing the human cerebral cortex on MRI scans into gyral based regions of interest. Neuroimage 31, 968–980.

Ehrlich, S., Brauns, S., Yendiki, A., Ho, B.C., Calhoun, V., Schulz, S.C., Gollub, R. L., Sponheim, S. R. (2012). Associations of Cortical Thickness and Cognition in Patients with Schizophrenia and Healthy Controls, Schizophrenia Bulletin, Volume 38, Issue 5, 1, Pages 1050–1062, https://doi.org/10.1093/schbul/sbr018

Fischl, B. and Dale, A.M., (2000). Measuring the Thickness of the Human Cerebral Cortex from Magnetic Resonance Images. Proceedings of the National Academy of Sciences, 97:11044–11049.

Fischl, B., Liu, A., Dale, A.M., 2001. Automated manifold surgery: constructing geometrically accurate and topologically correct models of the human cerebral cortex. IEEE Trans Med Imaging, 20(1):70–80.

Fischl, B., Salat, D.H., Busa, E., Albert, M., Dieterich, M., Haselgrove, C., van der Kouwe, A., Killiany, R., Kennedy, D., Klaveness, S., Montillo, A., Makris, N., Rosen, B., Dale, A.M., 2002. Whole brain segmentation: automated labeling of neuroanatomical structures in the human brain. Neuron 33, 341–355.

Fischl, B., van der Kouwe, A., Destrieux, C., Halgren, E., Segonne, F., Salat, D.H., Busa, E., Seidman, L.J., Goldstein, J., Kennedy, D., Caviness, V., Makris, N., Rosen, B., Dale, A.M., (2004b). Automatically parcellating the human cerebral cortex, Cereb Cortex 14, 11–22.

Fischl, B., Salat, D.H., van der Kouwe, A.J., Makris, N., Segonne, F., Quinn, B.T., Dale, A.M., (2004a). Sequence-independent segmentation of magnetic resonance images. NeuroImage 23 Suppl 1, S69–84.

Fischl, B. (2012). FreeSurfer. NeuroImage, 62(2), 774–781. http://doi.org/10.1016/j.neuroimage.2012.01.021

Fischl, B., Sereno, M.I., Tootell, R.B., Dale, A.M., 1999b. High-resolution intersubject averaging and a coordinate system for the cortical surface. Hum Brain Mapp 8, 272–284.

Fischl, B., Sereno, M.I., Dale, A.M., 1999a. Cortical surface-based analysis. II: Inflation, flattening, and a surface-based coordinate system. Neuroimage 9, 195–207.

Gur RE, Turetsky BI, Bilker WB, Gur RC (1999). Reduced gray matter volume in Schizophrenia, Arch Gen Psychiatry. 56(10):905–11.

Fusar-Poli, P., Radua, J., McGuire, P., Borgwardt S. (2012). Neuroanatomical Maps of Psychosis Onset: Voxel-wise Meta-Analysis of Antipsychotic-Naive VBM Studies, Schizophrenia Bulletin, Volume 38, Issue 6, 1, Pages 1297–1307, https://doi.org/10.1093/schbul/sbr134

Hooley J.M. (2010). Social Factors in Schizophrenia, Current Directions in Psychological Science, Vol 19, Issue 4, pp. 238–242, https://doi.org/10.1177/0963721410377597

Hofner B. & Hothorn T. (2017). stabs: Stability Selection with Error Control, R package version 0.6-3, https://CRAN.R-project.org/package=stabs.

Hofner, B., Boccuto L., and Goeker M. (2015). Controlling false discoveries in high-dimensional situations: Boosting with stability selection. BMC Bioinformatics, 16:144.doi:10.1186/s12859-015-0575-3>

Hotelling, H. (1933). Analysis of a complex of statistical variables into principal components. Journal of Educational Psychology, 24, 417–441, and 498–520.

Jovicich, J., Czanner, S., Greve, D., Haley, E., van der Kouwe, A., Gollub, R., Kennedy, D., Schmitt, F., Brown, G., Macfall, J., Fischl, B., Dale, A., (2006). Reliability in multi-site structural MRI studies: effects of gradient non-linearity correction on phantom and human data. Neuroimage 30, 436–443.

Joyal, C.C., Laakso, M.P., Tiihonen, J., Syvälahti, E., Vilkman, H., Laakso, A., Alakare, B., Räkkolainen, V., Salokangas, R.K.R., and Hietala J. (2003). The amygdala and Schizophrenia: a volumetric magnetic resonance imaging study in first-episode, neuroleptic-naive patients, Biological Psychiatry, Volume 54, Issue 11, 1, Pages 1302–1304, DOI:https://doi.org/10.1016/S0006-3223(03)00597-3

Hor K. & Taylor M. (2010) Review: Suicide and schizophrenia: a systematic review of rates and risk factors, Journal of Psychopharmacology, Vol 24, Issue 4_suppl, pp. 81–90, https://doi.org/10.1177/1359786810385490

Kalus P., Slotboom, J., Gallinat J., Wiest R., Ozdoba C., Federspiel A., Strik W. K., Buri C., Schroth G., Kiefer C., (2005) The amygdala in Schizophrenia: a trimodal magnetic resonance imaging study, Neuroscience Letters, Volume 375, Issue 3, 3 March, Pages 151–156 DOI: https://doi.org/10.1016/j.neulet.2004.11.004

Koutsouleris N., Meisenzahl E.M., Davatzikos C., Bottlender R., Frodl T., Scheuerecker J., Schmitt G., Zetzsche T., Decker P., Reiser M., Möller H.J., Gaser C. (2009). Use of neuroanatomical pattern classification to identify subjects in at-risk mental states of psychosis and predict disease transition, Arch Gen Psychiatry, 66(7):700–12. doi: 10.1001/archgenpsychiatry.2009.62.

Kuperberg, G.R., Broome, M.R., McGuire, P.K., David, A.S., Eddy, M., Ozawa, F., Goff, D., West, W.C., Williams, S.C., van der Kouwe, A.J., Salat, D.H., Dale, A.M., Fischl, B., 2003. Regionally localized thinning of the cerebral cortex in Schizophrenia. Arch Gen Psychiatry 60, 878–888.

Long X., Chen L., Jiang C., Zhang L., Alzheimer’s Disease Neuroimaging Initiative (2017) Prediction and classification of Alzheimer disease based on quantification of MRI deformation. PLoS ONE 12(3): e0173372. https://doi.org/10.1371/journal.pone.0173372

Martinez-Cantin, R. (2014). BayesOpt: A Bayesian Optimization Library for Nonlinear Optimization, Experimental Design and Bandits. Journal of Machine Learning Research, 15:3735–3739

Mechelli, A., Price, C.J., Friston, K.J. and Ashburner, J. (2005). Voxel-based morphometry of the human brain: Methods and applications. Current Medical Imaging Reviews, pages 105–113.

Meinshausen N. and Buehlmann P. (2010), Stability selection. Journal of the Royal Statistical Society, Series B, 72, 417–473.

Modinos, G., & McGuire, P. (2015). The Prodromal Phase of Psychosis. Current Opinion in Neurobiology, 30, 100–105. DOI: 10.1016/j.conb.2014.11.003

Monteiro, J. M., Rao, A., Shawe-Taylor, J., & Mourão-Miranda, J. (2016). A multiple hold-out framework for Sparse Partial Least Squares. Journal of Neuroscience Methods, 271, 182–194. http://doi.org/10.1016/j.jneumeth.2016.06.011

Nichols, T. E. et al. (2016) Best practices in data analysis and sharing in neuroimaging using MRI. Preprint at bioRxiv http://dx.doi.org/10.1101/054262

Palaniyappan L., Marques T.R., Taylor H., Handley R., Mondelli V., Bonaccorso S., Giordano A., McQueen G., DiForti M., Simmons A., David A.S., Pariante C.M., Murray R.M., Dazzan P. (2013). Cortical Folding Defects as Markers of Poor Treatment Response in First-Episode Psychosis. JAMA Psychiatry, 70(10):1031–1040. doi:10.1001/jamapsychiatry.2013.203

Palaniyappan L., Marques T.R., Taylor H, Mondelli V., Reinders A. A. T. S., Bonaccorso S., Giordano A., DiForti M., Simmons A., David A.S., Pariante CM, Murray RM, Dazzan P. (2016). Globally Efficient Brain Organization and Treatment Response in Psychosis: A Connectomic Study of Gyrification, Schizophrenia Bulletin, Volume 42, Issue 6, 1, 1446–1456, https://doi.org/10.1093/schbul/sbw069

Palaniyappan L. and Liddle P.F. (2012). Aberrant cortical gyrification in Schizophrenia: a surface-based morphometry study, J Psychiatry Neurosci; 37(6): 399–406.

Palaniyappan, L., Pavan, M., Verghese, J., Thomas P. White, Peter F. Liddle (2011). Folding of the Prefrontal Cortex in Schizophrenia: Regional Differences in Gyrification, Biological Psychiatry, Volume 69, Issue 10, 974–979

Pearson, K. (1901). “On Lines and Planes of Closest Fit to Systems of Points in Space”. Philosophical Magazine. 2 (11): 559–572. doi:10.1080/14786440109462720.

Segonne F, Dale AM, Busa E, Glessner M, Salat D, Hahn HK, Fischl B (2004). A hybrid approach to the skull stripping problem in MRI. Neuroimage 22:1060–1075

Segonne, F., E. Grimson and B. Fischl, (2003). Topology Correction of Subcortical Segmentation MICCAI.

Segonne, F., Pacheco, J. Fischl, B. (2007). Geometrically Accurate Topology-Correction of Cortical Surfaces Using Nonseparating Loops, IEEE Transactions on Medical Imaging, 26(4): 518–529.

Segonne, F., Grimson, E., Fischl B. (2005). Genetic Algorithm for the Topology Correction of Cortical Surfaces, IPMI, 393–405.

Selemon L.D., Mrzljak J., Kleinman J.E., Herman M.M., Goldman-Rakic P.S. (2003). Regional Specificity in the Neuropathologic Substrates of Schizophrenia: A Morphometric Analysis of Broca’s Area 44 and Area 9. Arch Gen Psychiatry; 60(1):69–77. doi:10.1001/archpsyc.60.1.69

Shah R.D. and Samworth R.J. (2013). Variable selection with error control: another look at stability selection. Journal of the Royal Statistical Society, Series B, 75, 55–80.

Sled, J.G., Zijdenbos, A.P., Evans, A.C., (1998). A nonparametric method for automatic correction of intensity nonuniformity in MRI data. IEEE Trans Med Imaging 17, 87–97.

Smith, S., & Nichols, T. (2017). Statistical Challenges in “Big Data” Human Neuroimaging, Neuron, Volume 9, Issue 2, 263–268

Tamminga, C.A., & Medoff, D.R. (2000). The biology of schizophrenia. Dialogues in Clinical Neuroscience, 2(4), 339–348.

Taylor, J.A., Matthews, N., Michie, P.T., Rosa, M.J., Garrido M.I. (2017). Auditory prediction errors as individual biomarkers of schizophrenia, NeuroImage: Clinical, 15, 264–273.

Tibshirani, R. (1996). Regression Shrinkage and Selection via the Lasso. Journal of the Royal Statistical Society. Series B (Methodological), 58(1), 267–288. http://www.jstor.org/stable/2346178

Varoquaux G. (in press). Cross-validation failure: small sample sizes lead to large error bars NeuroImage

Viher, P. V., Stegmayer, K., Kubicki, M., Karmacharya, S., Lyall, A. E., Federspiel, A., & Walther, S. (2018). The cortical signature of impaired gesturing: Findings from Schizophrenia. NeuroImage: Clinical, 17, 213.

Wagstyl, K., Ronan, L., Whitaker, K. J., Goodyer, I. M., Roberts, N., Crow, T. J. & Fletcher, P. C. (2016). Multiple markers of cortical morphology reveal evidence of supragranular thinning in schizophrenia, Translational Psychiatry, volume 6, page e780

Woo C.W., Chang L.J., Lindquist M.A., Wager T.D. (2017). Building better biomarkers: brain models in translational neuroimaging. Nat Neurosci. 23;20(3):365–377. doi: 10.1038/nn.4478.

Yeo, B.T., Sabuncu, M.R., Vercauteren, T., Holt, D., Amunts, K., Zilles, K., Golland, P., Fischl, B. (2010b). Learning Task-Optimal Registration Cost Functions for Localizing Cytoarchitecture and Function in the Cerebral Cortex, IEEE Transactions on Medical Imaging, 29(7):1424–1441

Yeo, B.T., Sabuncu, M. R., Vercauteren, T., Ayache, N., Fischl, B., Golland P. (2010). Spherical Demons: Fast Diffeomorphic Landmark-Free Surface Registration, IEEE Transactions on Medical Imaging. 29(3):650–668, 2010a

Zou, H. & Hastie, T. (2005). “Regularization and Variable Selection via the Elastic Net”. Journal of the Royal Statistical Society, Series B: 301–320.

